# Systematic screens for fertility genes essential for malaria parasite transmission reveal conserved aspects of sex in a divergent eukaryote

**DOI:** 10.1101/2023.12.25.572958

**Authors:** Claire Sayers, Vikash Pandey, Arjun Balakrishnan, Katharine Michie, Dennis Svedberg, Mirjam Hunziker, Mercedes Pardo Calvo, Jyoti Choudhary, Ronnie Berntsson, Oliver Billker

## Abstract

Sexual reproduction in malaria parasites is essential for their transmission to mosquitoes. It also offers a divergent eukaryote model to understand the evolution of sex. Through a panel of genetic screens, where each sex of *Plasmodium berghei* was mutagenised separately with barcoded vectors, we identify 401 sex and transmission-related gene functions and define roles for hundreds of unstudied fertility genes as putative targets for transmission blocking interventions. The functional data provide a deeper understanding of female metabolic reprogramming, meiosis and the axoneme. We identify a protein complex of a SUN domain protein, SUN1, and a moonlighting putative allantoicase, ALLC1, that is essential for male fertility by linking the microtubule organising centre to the nuclear envelope and enabling mitotic spindle formation during male gametogenesis. Both proteins have orthologs in mouse testis, and the data point to an ancient role for atypical SUN domain proteins in fertility. Altogether, our data provide an unbiased picture of the molecular processes that underpin malaria parasite transmission but also highlight ancestral aspects of sex that have evolved close to the last eukaryotic common ancestor.

## INTRODUCTION

The ancestral eukaryote introduced a significant innovation in genetic recombination through sexual reproduction. Sex evolved in the context of a life cycle alternating between haploid and diploid phases, in which flagellated haploid cells differentiate into gametes, which subsequently fuse to create a diploid zygote undergoing meiosis^1^. Our current understanding of the molecular mechanisms governing sexual reproduction primarily stems from research in model organisms. For instance, the yeast *Saccharomyces cerevisiae* reproduces asexually as diploids but can undergo meiosis to produce haploid gametes under stress. Genetic screens in this organism have identified the genes essential for meiosis^2^. In multicellular model species where sexual reproduction is obligatory, studying fertility encounters challenges, especially when sterile mutants are difficult to propagate. Consequently, limited genome-scale fertility studies are available for such organisms^3^. While saturated collections of *Arabidopsis thaliana* mutants have enabled the identification of mutants affecting sexual processes, intricate and labour-intensive screens are necessary to distinguish between fertilisation and gametophyte developmental phenotypes^5,7^. Similarly, in *Drosophila melanogaster*, a collection of sterile mutants has been utilised to explore meiosis and gamete development^9^. Notably, these studies are limited to a small portion of the eukaryotic tree of life, specifically the Opisthokonta and Chloroplastida clades, accounting for only a fraction of eukaryotic diversity^11^. In contrast, malaria parasites, belonging to the alveolates, represent a divergent clade that, along with stramenopiles and Rhizaria, comprises nearly half of all eukaryotic species diversity^13^. Systematic exploration of sexual processes within this diverse clade is crucial for uncovering conserved molecular mechanisms of sexual reproduction that trace back to the ancestral eukaryote. One such example is the conserved gamete fusion protein hapless-2 (hap2), whose function in eukaryotic gamete fusion was elucidated through comparative studies in malaria parasites, plants, and green algae^15,16^. Interestingly, this protein has been found to share a common origin with certain viral fusogens^18^.

Understanding sex in *Plasmodium* is of particular importance because malaria remains a global health concern. An estimated 608,000 people died from the disease in 2022, most of them children under the age of five (WHO World malaria report, 2023). Haploid parasites reproducing asexually inside erythrocytes are alone responsible for causing disease, but sexual reproduction is essential for the parasite to infect the mosquitoes that serve as obligatory vectors. Polygenomic (multi-strain) infections are common in high transmission areas^20^, where meiotic recombination between strains generates and maintains genetic diversity in populations^21^. Sex contributes to the spread of drug resistance alleles, the evolution of virulence mechanisms and the diversification of a large repertoire of surface antigens that is key to immune evasion and has to date prevented the development of vaccines against the pathogenic blood stage of any *Plasmodium* species.

During each asexual round of replication, a subpopulation of intraerythrocytic blood stages leaves the asexual cycle (Fig. 1A) and transitions to male or female gametocytes that become infectious to mosquitoes when fully mature^22,23^. Once taken up into the mosquito midgut, these sexual precursors respond immediately to environmental triggers^24^, egress from their host cells, and differentiate into gametes, which fuse to form a zygote. The time-limiting factor in gamete formation is the threefold replication of the male genome and the cytosolic assembly of axonemes that precede the endomitotic formation of eight flagellated microgametes in a rapid process termed exflagellation which is completed within around 20 minutes. After fertilisation and nuclear envelope fusion, a premeiotic genome replication is followed by two rounds of meiosis. Without undergoing cytokinesis, the zygote transforms into a motile ookinete, which invades the midgut epithelium and forms a cyst. Meiosis produces four haploid genomes. These probably all persist within the same nuclear envelope, since the growing cyst expresses the genes inherited from both parents as it replicates by closed mitosis. Functionally, the ookinete and oocyst therefore behave like diploid cells. Their individual genomes only segregate during sporozoite formation approximately one to two weeks later. Haploid sporozoites egress from the cyst and invade the salivary glands of the mosquito, who delivers them into the skin of the next host. They make their way to the liver where they resume asexual replication, completing their life cycle.

**Figure 1.**
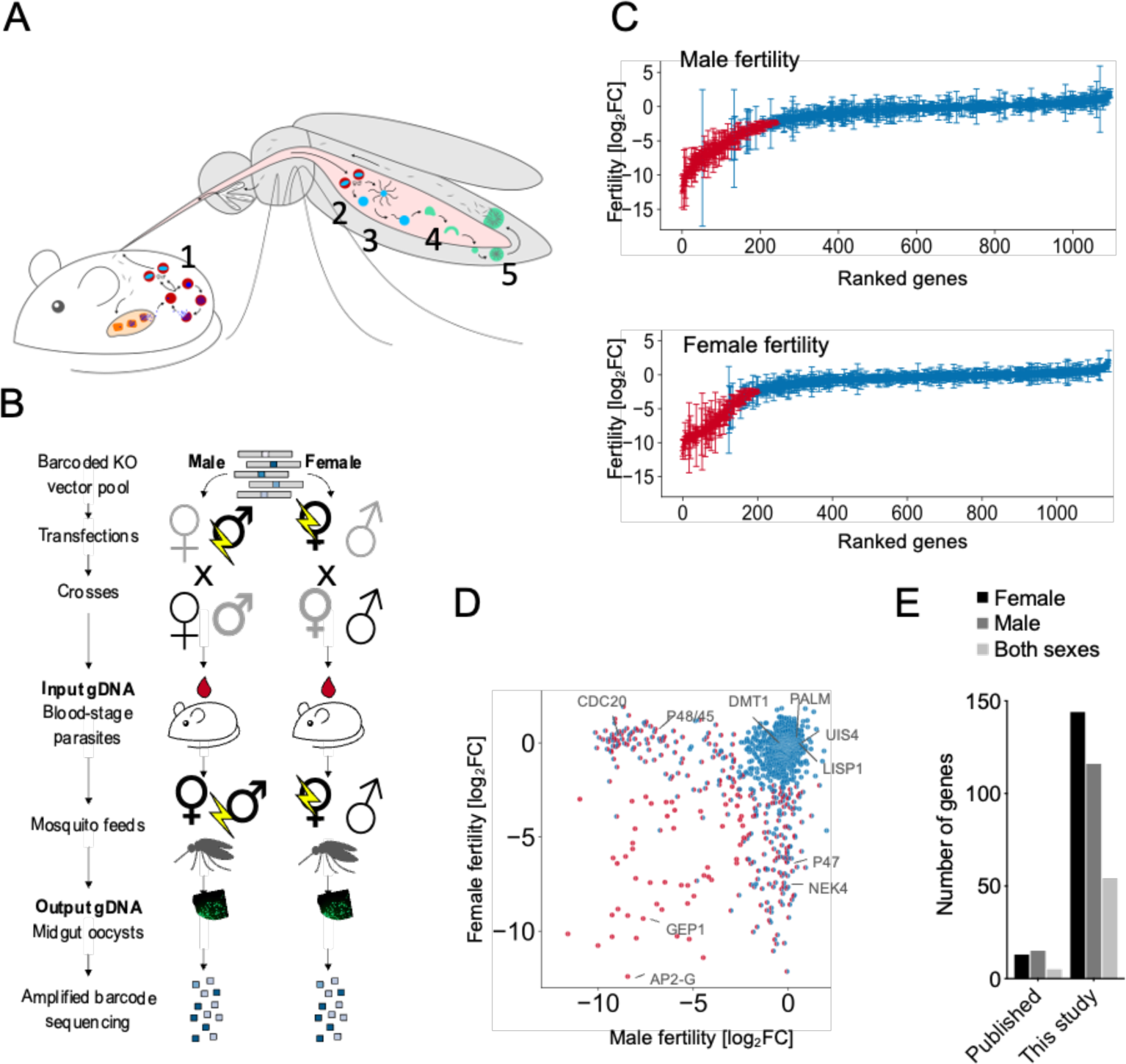
Sex-specific genetic screens identify hundreds of fertility genes in *P. berghei*. (A) Life cycle diagram showing biological events contributing to fertility: Gametocyte formation and maturation (1), gametocyte activation (2), fertilisation (3) and ookinete development (4). Endpoint of fertility screens is the oocyst (5). (B) Schematic illustrating the design of sex-specific knockout screens. (C) Female and male fertility plotted separately for each screen. Red indicates reduced fertility; blue data points are not significantly reduced. Error bars show 2x standard deviation. (D) Data as in (C), but male and female fertility are plotted against each other to reveal genes with dual functions. Colours as in (C), with the left semi-circle showing female fertility. Selected genes with published phenotypes are labelled. (E). Number of sex-specific fertility phenotypes from crosses in the current screen as compared to previously published data^31^.

Current antimalarials were developed to target the asexually replicating blood and liver stages of *Plasmodium*, but for the local elimination and global eradication of malaria, reducing parasite transmission by mosquitoes is now considered essential^25,26^. Among the candidates for transmission blocking targets are hundreds of genes and proteins whose abundance increases strongly during sexual development^12^. Few sex-specific genes have experimentally assigned functions, including leading antigens for transmission blocking vaccine development^27,28^ and a validated target for small molecules that block transmission^29^.

The cell biology of *Plasmodium* gametes and zygotes remains a major knowledge gap. We therefore set out here to identify *Plasmodium* sterility genes systematically, making use of the *Plasmo*GEM resource of barcoded gene knock-out (KO) vectors that allows the targeted disruption of more than two thirds of all protein-coding genes in the core genome of *Plasmodium berghei*, a parasite infecting rodents^19,30^. We first report the results of two signature-tagged mutagenesis screens designed to identify separately the sterile mutants in the male and female developmental lineages. We then stratify male sterility mutants further in a motility screen, and we integrate phenotype data with structure predictions and gene expression information to delineate the biological processes essential for the sexual development of a malaria parasite. We identify a new protein interaction between a moonlighting allantoicase and a conserved nuclear envelope protein that is important to couple the microtubule organising centre to the nucleus during microgamete formation. We present evidence suggesting the complex could be conserved in human testis, illustrating the potential of a global functional analysis in a divergent eukaryote to identify broadly conserved features of sexual development.

## RESULTS

### Genome-scale screens identify hundreds of new fertility genes

To screen for sterile *Plasmodium* mutants, we transfected pools of barcoded *Plasmo*GEM KO vectors separately into two mutant lines (Fig. S1), each capable of producing gametocytes of only the male or female sex. Mice were infected with pools of mutagenised parasites, and a complementary line was spiked in to donate wild type (WT) gametocytes of the missing sex. Fertility of the mutagenised sex was then assayed by allowing *Anopheles stephensi* mosquitoes to feed. After 13 to 14 days, mosquito midguts with oocysts were dissected and barcodes were counted by DNA sequencing to quantify transmission success of mutants from either the male or the female parent (Fig. 1B). Included in the screens were all genes with *Plasmo*GEM vectors and which were non-essential in asexual blood stages (growth rate of >0.5^12^). The population bottleneck during transmission made it necessary to study mutants in batches of 50-200 and combine data from hundreds of experiments and >3000 individually dissected midguts to assess 1302 genes for sex specific roles in parasite transmission.

At the chosen significance level, female fertility was reduced in 220 mutants and male fertility in 181 (Fig. 1C) relative to a set of control mutants in each pool. Most phenotypes were limited to one sex, but 54 mutations affected both (Fig. 1D). 130 hits were only known as “conserved *Plasmodium* proteins” but searching protein structure databases with AlphaFold predictions provided further functional annotation for 63 of these (Table S1). The screens increased by almost 10-fold the number of *P. berghei* genes for which sex-specific phenotypes are known from crosses (Fig. 1E). 89% of published phenotypes were reproduced (Table S1), discrepancies being mostly borderline cases, where variance was high or biological effects small.

### Few genes affect fertility of both sexes

Having evolved from an isogamous alveolate ancestor^32^, malaria parasites have become markedly anisogamous. *P. berghei* sexes therefore do not share many biological systems beyond general cell maintenance. Among the mutants affecting both sexes are known transcriptional regulators important for the initial commitment to sexual development and the execution of the sexual development programmes^33–35^. Posttranscriptional control is equally important for gametocytogenesis and zygote development^12,36,37^, and the fertility screens reveal over a dozen putative RNA binding and processing proteins (Table S3) that provide starting points to unravel the mechanisms underpinning gametocyte differentiation. Data also point to roles for protein ubiquitination and to regulation of translation through diphthamide modification, which typically is targeted to eukaryotic elongation factor 2 (Table S2), and which may contribute to enhance translation when gametocyte activation triggers the de-repression of stored mRNAs through an unknown mechanism^12,36^. Another shared system likely enables gametocytes to recognise the known triggers that signal arrival in the mosquito midgut^38^, which may include an elusive receptor for the mosquito catabolite xanthurenic acid known to serve as signal^24^ and downstream transduction proteins^39^.

### Many female fertility genes reflect changes in carbon metabolism

When mapped onto the co-expression clusters of the *P. berghei* single-cell transcriptomic atlas^10,16,17^ (Fig. 2A), female phenotypes were less enriched for genes with female-specific transcripts of clusters 13 and 14 than for the more ubiquitously transcribed genes in expression cluster 4. The latter account for a marked enrichment of a subset of mitochondrial metabolic pathways among all female fertility genes (Fig. 2B). The *Plasmodium* mitochondrion is an essential organelle^10,17,18^ and validated drug target of asexual blood stages, which the zygote inherits exclusively from the female gamete^40^. This is consistent with the uniparental inheritance mode that is common among eukaryotes and one of the hallmarks of sex. Female fertility genes now provide a detailed view of the metabolic reprogramming that accompanies transition to the vector (Fig. 2C, D; Table S2).

**Figure 2.**
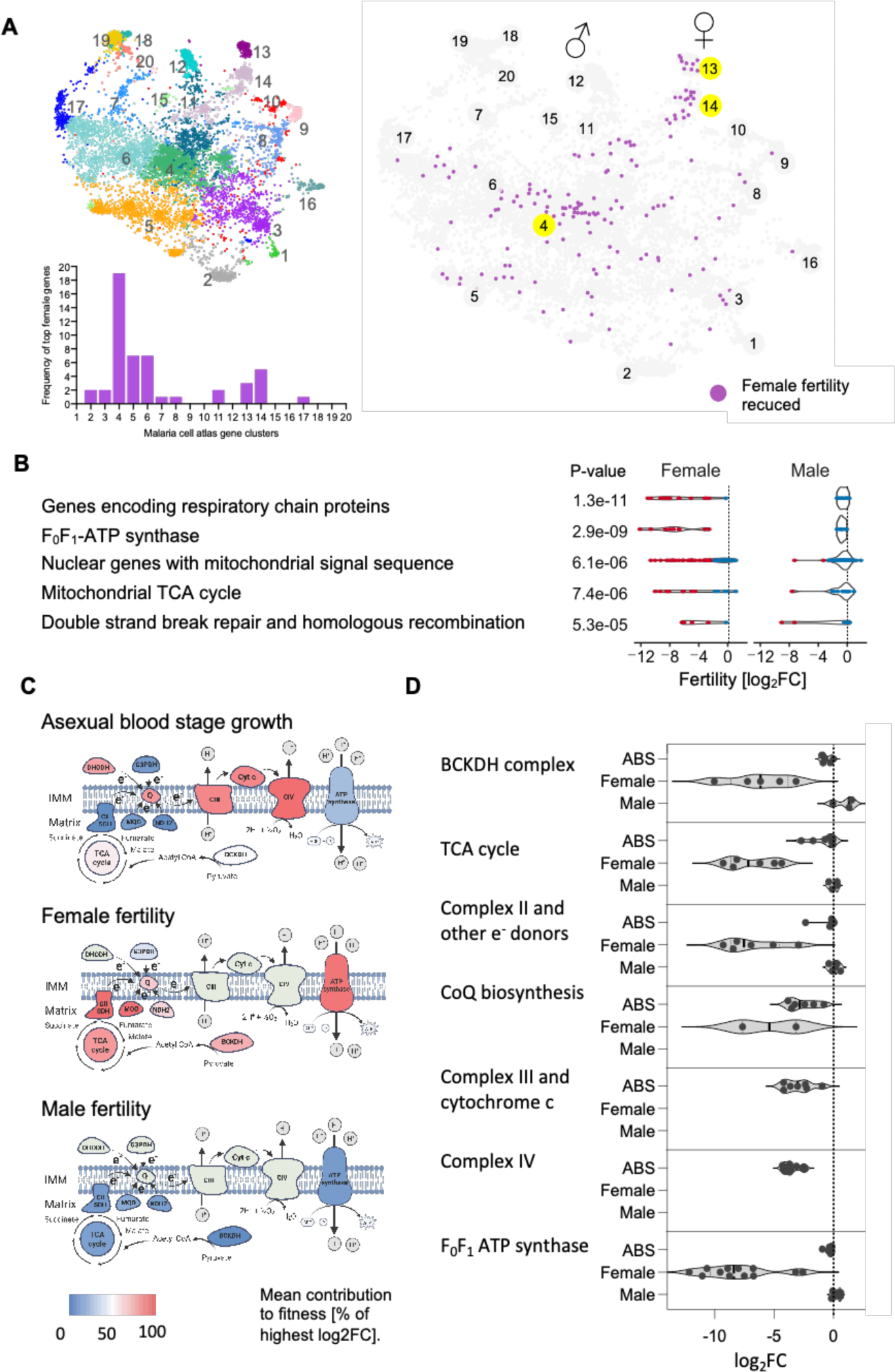
Female fertility genes and the role of mitochondrial metabolic reprogramming. (A) Gene expression characteristics of female fertility genes visualised using a k-nearest neighbour graph of gene expression profiles (numbered) from the malaria cell atlas^21^. Bar chart shows the distribution of the top 50 screen hits over gene expression profiles. (B) Enrichment analysis of curated gene functions from the Malaria Parasite Metabolic Pathways database^43^. (C) Graphic representation of the mitochondrial electron transport chain comparing the relative importance of metabolic subsystems and protein complexes in asexual blood stage fitness^27^ and in female and male fertility as the arithmetic mean of all tested components and normalised to account for the unique dynamic range of each screen. DHODH dihydroorotate dehydrogenase, G3DPH, Q, CII SDH, MQO, NDH2 - glycerol-3-phosphate dehydrogenase, TCA - tricarboxylic acid, BCKDH, Q, CIII - complex III, CIV - complex IV, Cyt c - cytochrome c. (D) Violin plots showing phenotypes of individual genes tested from each biological subsystem. Abbreviations as in C. Note that the apparent smaller effect size of asexual fitness effects is due to the smaller dynamic range inherent to that type of screen. Data for individual genes are shown in Table S2.

Although the core of the mitochondrial electron transport chain (mETC), from the initial electron acceptor coenzyme Q with its biosynthetic pathway to the proton-pumping complexes III and IV, cannot be disrupted in asexual blood stages^27^ (Fig. 2C, D; Table S2), its main function is not in energy production since parasites in the glucose-rich environment of the blood rely largely on cytosolic glycolysis^41^. Central carbon metabolism is remodelled in mosquito stages, where an intact tricarboxylic acid (TCA) cycle and oxidative phosphorylation^42^ must be inherited from the female gamete (Fig. 2C, D). The validated antimalarial target dihydroorotate dehydrogenase, which also functions in pyrimidine biosynthesis, is the only blood-stage essential electron donor to coenzyme Q, but three other single-subunit dehydrogenases, as well as complex II of the electron transport chain, become individually essential for female fertility (Fig. 2C, D), although all are expressed throughout the life cycle, and some have been pursued as drug targets in *P. falciparum*.

Since the main task of mitochondrial electron transport is not in energy production, subunits of the F^0^-F^1^ ATP synthase complex, which couples the proton-motive force generated by proton pumping to ATP production, contribute little to *P. berghei* asexual erythrocytic growth, but only become essential for female fertility (Fig. 2C, D). This corroborates an earlier finding that the ATP synthase β subunit is important for zygotes to form and transform into ookinetes in the midgut^44^. In the related apicomplexan *T. gondii* one of the recently discovered non-canonical ATP synthase components is needed to shape the inner mitochondrial membrane into cristae^45–47^, a process which in *Plasmodium* occurs during sexual differentiation of gametocytes^44,47,48^. We found five *P. berghei* homologues of non-canonical ATP synthase components that contribute to female fertility (Table S2), and which may serve as starting points for understanding how mitochondrial function relates to the formation of protein supercomplexes and cristae^49^.

Non-mitochondrial female fertility genes often show female-specific expression and are enriched for DNA recombinases (Fig. 2B), which may function in homologous recombination during meiosis. *Plasmodium* parasites have been underutilised as a model to study meiosis in a divergent eukaryote^50^. Disrupting the meiotic recombinase *dmc1* (disrupted meiotic cDNA 1) resulted in a significant drop in female fertility, recapitulating a previously reported block in oocyst sporogony of a *dmc1* mutant^51^. Other female fertility genes that could be part of the homologous recombination machinery include *rad50* (PBANKA_0104600), an AlphaFold hit to *rad51* (PBANKA_0939500), and a *rad54*-like putative DNA helicases (PBANKA_1226800) (Table S2). Other meiotic recombination genes of *P. berghei* did not yield data in the screens, including *spo11* homologues with predicted roles in inducing meiotic double strand breaks, and the meiotic nuclear division protein 1, *mnd1* (PBANKA_1325200).

A synaptonemal complex has been seen by transmission electron microscopy between homologous chromosomes^52^ in the *Plasmodium* zygote but its molecular composition remains unknown. HORMA domain proteins, Hop1 and Hop2, and a recently identified divergent homolog of synaptonemal complex protein 2^53^ are possible constituents. *Hop1* (PBANKA_1407900) and *sycp2* (PBANKA_1026300) are both expressed most highly in female gametocytes and dispensable for asexual growth, but the screens show no loss of fertility, suggesting failure to undergo synapsis may be less catastrophic for oocyst development than failure to resolve double-strand breaks in an organism that lacks non-homologous end joining-mediated repair. We do, however, note a moderate but significant reduction in female fertility associated with disrupting PBANKA_1419300, a protein for which we predict a Rec21-Rec8 domain, and which could therefore be a homolog of the meiosis-specific cohesin Rec8.

### A motility subscreen characterises male fertility genes

Male fertility phenotypes are highly enriched for male-specific expression clusters 11 and 12 of the malaria cell atlas (p = 9.8e-13 and 1.2e-12, respectively; Fig. 3A), and the most enriched functional gene categories relate to mitosis, through which microgametes are produced, and to components of the flagellum, which powers microgamete movement (Fig. 3B). We stratified the top 125 male genes further by applying a two-step centrifugation protocol to a pool of barcoded mutants^54^. In the first step egressed gametes were enriched based on their low density. In the second step, motile cells were allowed to swim up from a pellet of egressed gametes to enrich for mutants in which motility was not compromised (Fig. 3C). We predicted that mutants which according to a gametocyte development screen^34^ failed to establish the full transcriptional program of mature microgametocytes would likely be blocked before egress. Confirming this notion, a reduced egress score (Fig. 3D; Table S3) was highly correlated with a male gametocyte developmental phenotype (*X^2^* (1, *N* = 108) = 18.77, *p* = *0.000015*), while a low motility score (Fig. 2E; Table S3) was not (*X^2^* (1, *N* = 108) = 0.288, *p* = *0.62*), validating the secondary screens for the further stratification of male sterility genes.

**Figure 3.**
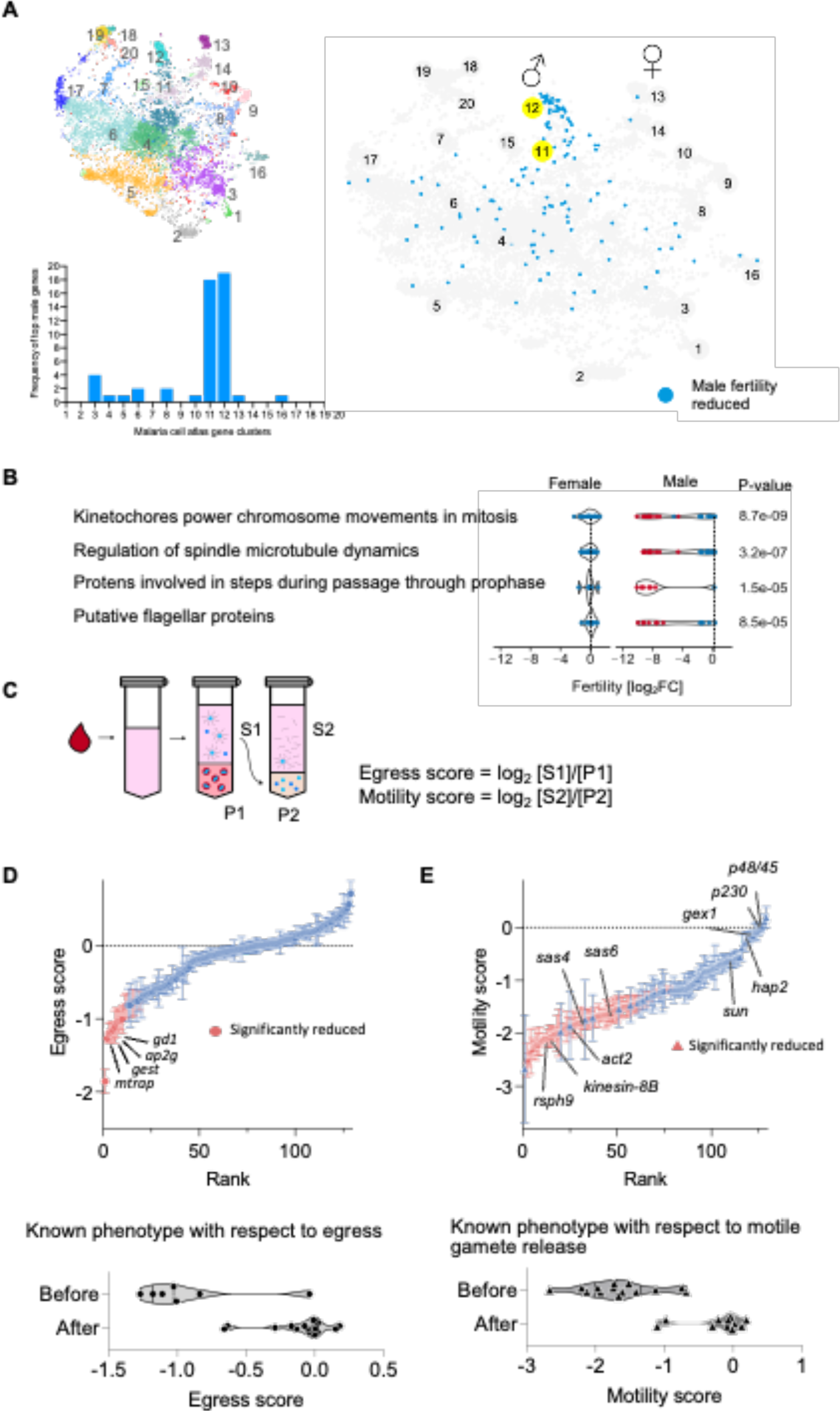
Subscreens of male fertility genes. (A) Gene expression characteristics of male fertility genes visualised using a k-nearest neighbour graph of gene expression profiles (numbered) from the malaria cell atlas^21^. Bar chart shows the distribution of the top 50 screen hits over gene expression profiles. (B) Enrichment analysis of curated gene functions from the Malaria Parasite Metabolic Pathways database^43^. (C) Schematic representation of a secondary screen for 125 male fertility mutants for developmental blocks before egress and before microgametes become motile. (D) Mutants ranked by egress score (top) and observed vs. expected phenotypes (bottom). (E) Mutants ranked by motility scores (top) and observed vs. expected phenotypes (bottom).

Malaria parasites have a typical flagellum with two central single microtubules connected to nine outer microtubule doublets nucleated from a basal body. It is used only for reproduction, has a short working life of about an hour and is assembled rapidly in the cytosol, presumably obviating the need for intraflagellar transport. A total of 24 genes in the screens were predicted by sequence and structural homology (Table S1) to be part of the axoneme, and many of these are enriched in the microgamete proteome^54^. 21 have a role in male fertility (Fig. 4A; Table S2), and all have normal egress but lower motility scores (Fig. 4B; Table S2), as would be expected of flagellar proteins.

**Figure 4.**
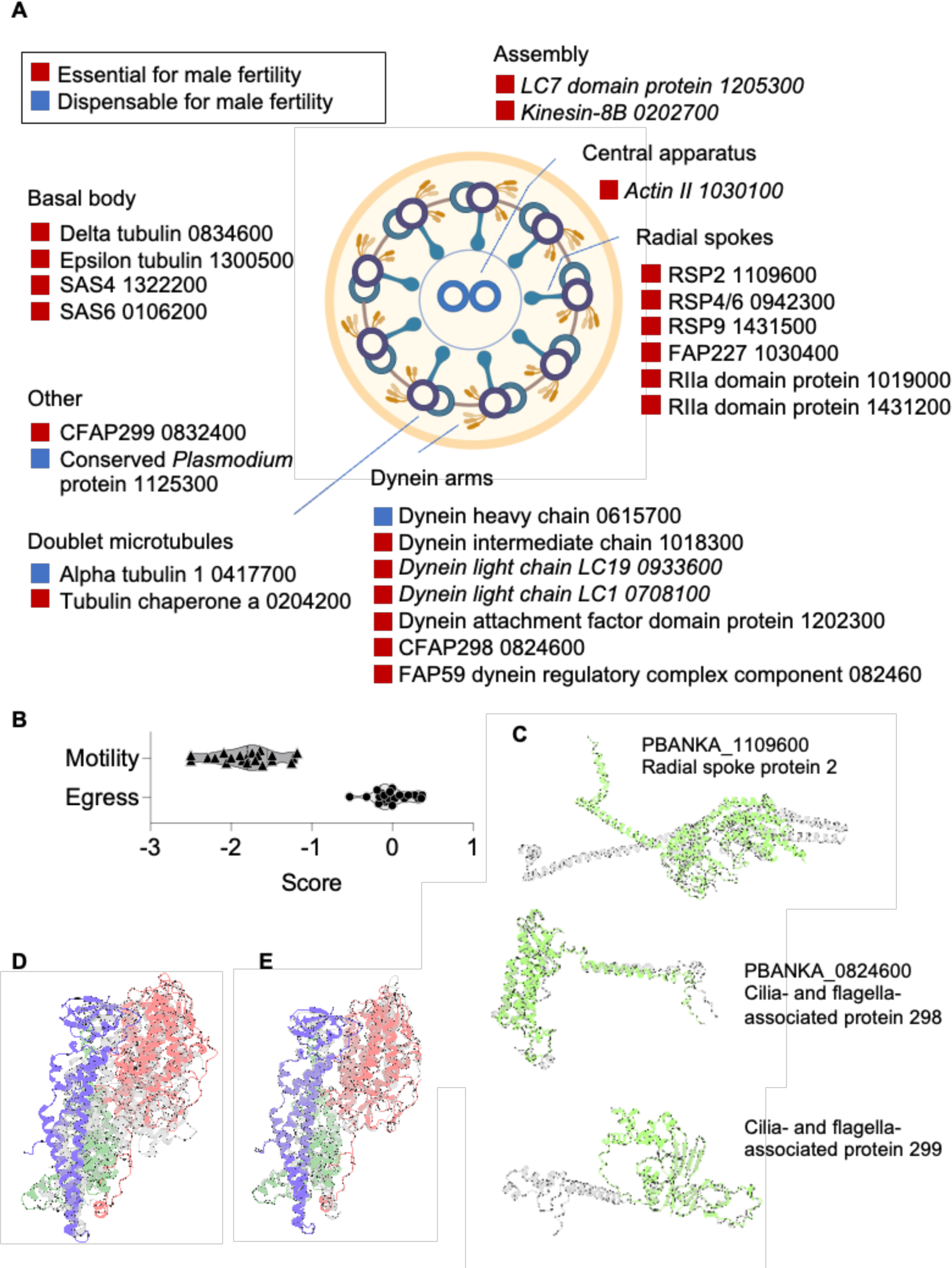
Male fertility genes. (A) A schematic of the axoneme with putative genes encoding its components. Numbers are the numerical part of the PBANKA gene IDs. (B) Motility and egress scores of putative axoneme components with sterility phenotype. (C) AlphaFold models of selected undescribed axoneme components in green, superimposed to their homologous human models in grey (PDB Code 7JTK from *Chlamydomonas reinhardtii* and AlphaFold entries A0A653H2U9 and A0A1D3CW47 (clockwise from top). (D) Structure model of the generic *P. berghei* GINS complex (Psf2 - red, Psf3 – green, generic combined Sld5-Psf1 - blue), superimposed on the corresponding subunits from the *H. sapiens* replisome complex in grey (PDB Code: 7PLO, chains D, E, F & G). (E) Generic GINS complex (coloured) on the gametocyte-specific GINS complex (grey), both from *P. berghei*. The structural models are highly similar, with rmsd values between 1-2 Å for the different subunits. See Fig. S2 for gene IDs and further validation data.

How far components of the *Plasmodium* flagellum have diverged from model organisms is illustrated by the sparsity of known radial spoke proteins (RSPs), which are required to generate a flagellar beat. The current list includes orthologs of RSP9, which is required for motile axonemes, RSP3 and RSP4/6^55^. We additionally predict an RSP2 ortholog among the male sterility genes, a homologue of flagellar-associated protein 227 and two additional proteins with RIIa domains typically found in RSPs (Table S2), whose participation in the axoneme is less certain. Altogether, AlphaFold structure predictions reveal eight previously unidentified putative flagellar components among the male fertility genes, including proteins of the dynein regulatory complex (Fig. 4C; Table S2).

Simultaneously with axoneme assembly, microgametocytes replicate their genomes three times in under 10 minutes, requiring an estimated 1300 origins if the replication fork moves at normal speed^56^. It is unknown how replication in microgametocyte differs from asexual blood stages, whether replication origins are indeed amplified and how replication stops after three rounds in the absence of cell cycle checkpoints^57^. CDC45 and other canonical components of replication complexes are unsurprisingly essential in the blood stages (Table S2), but structural modelling of male fertility genes reveals starting points for answering these questions (Table S2), namely an alternative origin recognition complex 3 (ORC3) protein, a structural paralog of subunit 5 of the replication factor C (RFC) complex, a putative RAD54 recombinase, and two putative members of the heterotetrameric go-ichi-ni-san (GINS) complex, which is essential for the establishment of DNA replication forks and replisome progression^58^. Structural annotation revealed that *P. berghei* GINS is unusual in that two subunits of the canonical eukaryotic complex are fused into a single polypeptide, while the remaining two are each duplicated, with one paralog of each expressed most highly in male gametocytes and essential specifically for male fertility (Fig. 4D). The entire GINS complex can be modelled equally well with the male-specific and the generic components (Fig. S2), suggesting the complex may exist in different forms. Male-specific versions of GINS and possibly of ORC, MCM and RFC complexes may be important for some of the unique aspects of replication in male gametocytes.

Most male fertility genes function before gamete motility, but a small group of 22 motile, yet sterile mutants highlighted in Figure 4B and listed in Table S7 are of particular interest, since this group contains known surface-exposed microgamete proteins that are targets for transmission-blocking antibodies, such as HAP2, P230 and P48/45. Studies of other genes in this group will likely identify urgently needed new vaccine targets^59^, with candidates including several unannotated genes with predicted signal peptides, such as PBANKA_1329000, which is specifically expressed in male and contains two transmembrane domains (Table S2), suggesting it could be an unstudied essential microgamete surface protein.

### SUN1 is a nuclear envelope protein that connects axonemes, mitotic spindles and microgamete DNA

Among the male fertility mutants with normal motility scores is a protein with a carboxy-terminal SUN (<u>S</u>ad1p/<u>UN</u>C-84) domain, PBANKA_1430900, which we term SUN1 (Fig. 4B; Table S7). SUN domain proteins were present in the first eukaryote and typically span the inner nuclear envelope, where they form part of the LINC (linker of nucleoskeleton and cytoskeleton) complex that anchors the nucleus by connecting the nuclear lamina to the cytosolic cytoskeleton^60^. The lack of nuclear lamins and canonical LINC complex components in apicomplexa^61^ led us to hypothesise that SUN1 has a non-canonical function in male gamete formation. A *sun1* KO clone (Fig. S5) recapitulated the screen data. Consistent with their high mobility score in the sub-screen, *sun1* KO microgametocytes formed exflagellation centres when activated *in vitro* at the same rate as WT (Fig. 5A). Morphologically mature ookinetes were formed (Fig. 5B), but very few oocysts were present on the midguts of infected mosquitoes (Fig. 5C; Table S8). *sun1* KO ookinetes contained only half the DNA of WT (Fig. 5D), probably because most microgametes failed to deliver DNA to the zygote (Fig. 5E). This was not the result of defective replication during gametogenesis (Fig. 5F), but mitotic spindles were missing from nuclei (Fig. 5G), suggesting SUN1 is important for the microgamete nucleus or its genome to connect to the axoneme.

**Figure 5.**
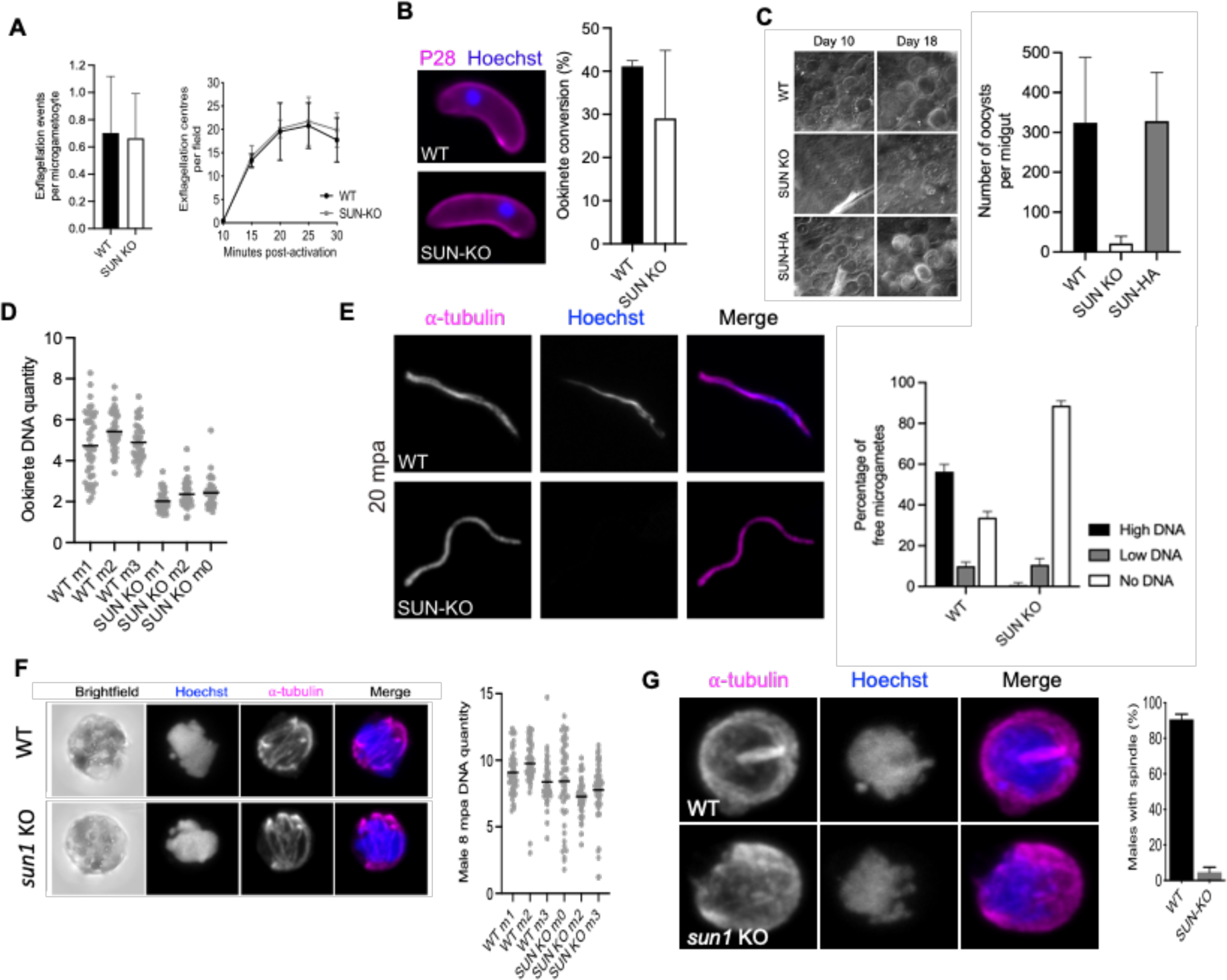
Characterisation of a *sun1* KO mutant. (A) Relative exflagellation rates and exflagellation time courses as determined by observing whole blood under Vaseline rimmed coverslips by phase contrast microscopy. (B) Fluorescence micrographs of ookinetes and ookinete conversion rates determined in whole blood cultured for 24 hours after gametocyte activation and stained with Cy3 conjugated mouse monoclonal antibody 13.1 against the zygote/ookinete surface protein P28. Conversion rates are expressed as the percentage of P28 positive cells that have transformed from a round stage to the typical ookinete shape. (C) Differential interference micrographs of infected midguts and oocyst counts on midguts dissected 10 and 18 days after feeding on infected mice. At least 16 midguts per mosquito were inspected per condition and replicate. (D) Hoechst fluorescence of ookinete nuclei measured relative to ring stage parasites containing 1N DNA. (E) Confocal fluorescence micrographs of representative microgametes and bar chart showing DNA content as judged by Hoechst fluorescence. (F) Representative micrographs of microgametocytes eight minutes after activation, when replication is usually complete and relative DNA content as determined above. (G) Representative micrographs of microgametocytes two minutes post-activation to illustrate the spindle of mitosis I in the nucleus of the WT cell and a bar chart showing relative abundance of microgametocytes with spindle. All bar charts and line diagrams show means of at least three replicates per line. Scale bars are 2 μm.

A SUN1 protein with a C-terminal haemagglutinin (HA)-epitope tag expressed from its endogenous locus (Fig. S4) did not interfere with protein function as judged by the production of normal numbers of oocysts by the tagged line (Fig. 5C). SUN1-HA was not detected in asexual blood stages and female gametocytes (not shown) but present in male gametocytes (Fig. 6 A-J). Cellular events of microgametogenesis are well characterised by ultrastructure expansion microscopy (U-ExM) and staining for α-tubulin, the centrosome marker centrin, the general protein label *N*-hydroxysuccinimide (NHS)-ester highlights microtubule organising centres (MTOCs) and the axonemes and spindles that emanate from them^60^. SUN1-HA was associated with the nuclear envelope of expanded microgametocytes (Fig. 6A), and two minutes after activation some of the protein formed two rings, one at the centre of each windmill-like tetrad of budding axonemes that were present at the opposite poles of the first endomitotic spindle (Fig. 6A). Labelling centrin to mark the cytosolic part of the MTOCs corroborated the nuclear envelope localisation of SUN1-HA (Fig. 6B). In expanded *sun1* KO gametocytes the two tetrads of forming axonemes were positioned on top of each other, consistent with the absence of a mitotic spindle that would be required for their separation along the nuclear envelope (Fig. 6C). Often these structures appeared in the cytosol distant from the nuclear envelope (Fig. 6D), suggesting SUN1 may be required to capture the MTOCs onto the nuclear membrane (Fig. 6E).

**Figure 6.**
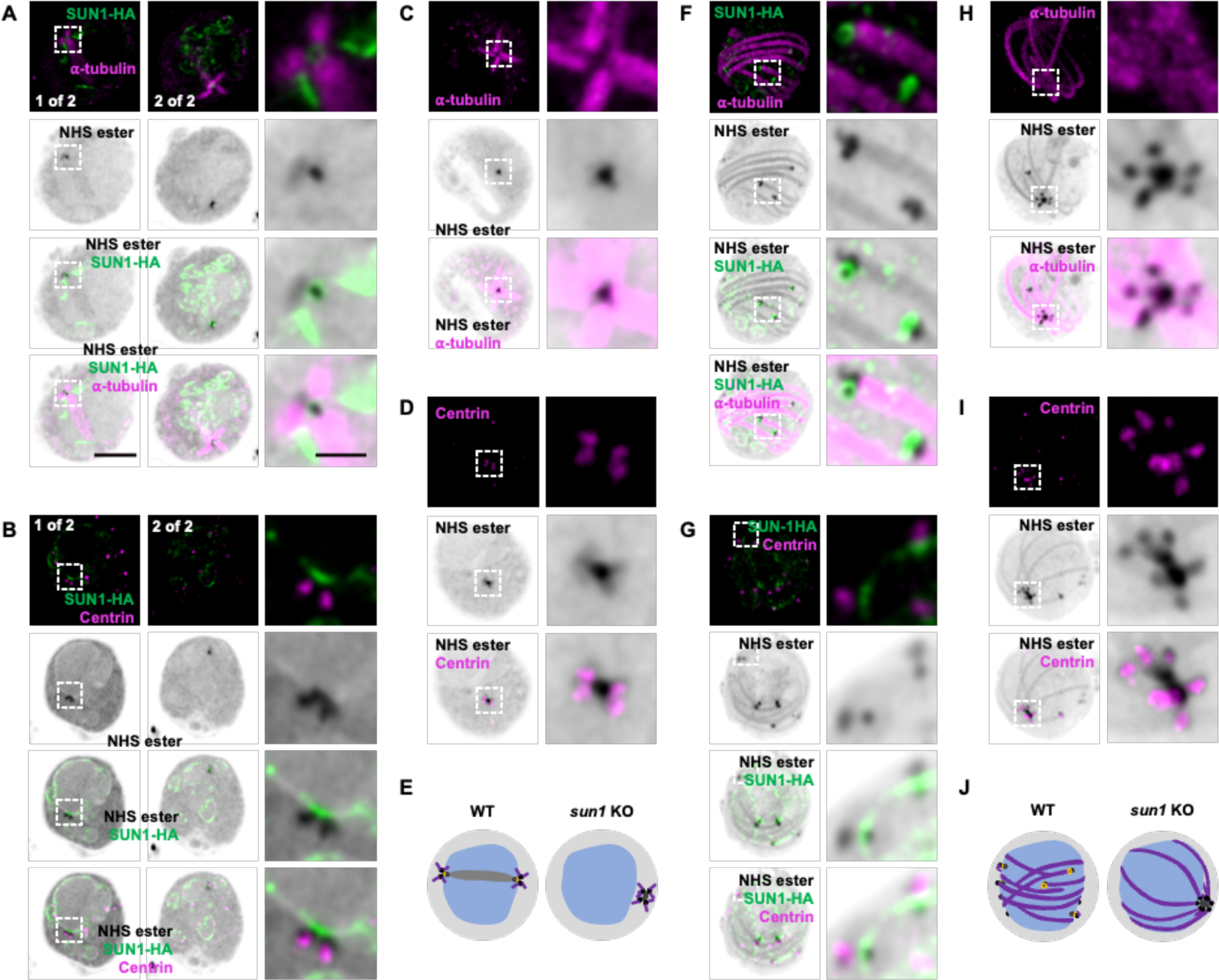
Ultrastructure expansion microscopy of microgametocytes lacking SUN1 or expressing SUN1-HA. Fluorescence detection of different proteins in individual confocal sections. All cells were also stained with Atto 594 NHS ester as a general protein label. Scale bars represent 5 μm of the expanded cell, 1 μm for the zoomed inserts. (A) A microgametocyte 2 minutes after activation, with SUN1-HA and α-tubulin labelled in two different confocal sections, each showing an axoneme tetrad emerging from the MTOCs on each end of the first mitotic spindle. (B) Single MTOC of a SUN1-HA microgametocyte 2 minutes post-activation and labelled as in A. (C, D) Confocal sections through the single MTOC of *sun1*-KO microgametocyte 2 minutes after activation, stained as indicated. E. Model of microgametocytes 2 minutes after activation. Nucleus blue, MTOCs black, emerging axonemes purple, SUN1 yellow. (F, G) Representative confocal section through an expanded microgametocyte expressing SUN1-HA eight minutes after activation, when axonemes are fully developed and three rounds of mitosis have been completed. (H, I) Eight minutes post-activation *sun1*-KO microgametocytes. (J) Model of microgametocytes eight minutes after activation. Colours as in (E).

Eight minutes after activation, when WT microgametocytes have completed their third endomitosis, the base of each axoneme was marked by a SUN1-HA ring (Fig. 6F) located between the cytosolic and the nucleoplasmic portion of each bipartite MTOC (Fig. 6G), precisely where spindle microtubules would need to pass through a nuclear pore to connect a haploid set of chromosomes to the centriolar region that has given rise to the axoneme (Fig. 6H). At the same time point, *sun1* KO microgametocytes also had elongated axonemes, which were however, emanating from MTOCs that remained connected, consistent with an absence of the mitotic events which in WT split the eight MTOCs and distribute them around the nuclear envelope (Fig. 6H-J).

### SUN1 interacts with an allantoicase-like protein

Immunoprecipitation of SUN1-HA from gametocyte lysates eight minutes post-activation identified one of two allantoicase-like proteins encoded in the *P. berghei* genome (Fig. 7A; Table S4), which we designated ALLC1. A reciprocal pull-down of ALLC1-HA (Fig. S4; Table S4) confirmed the interaction with SUN1 (Fig. 7A). Both proteins also interacted with the essential endoplasmic reticulum chaperone BiP and a DDRGK-motif containing conserved *Plasmodium* protein expressed most abundantly in male gametocytes. While the disruption of the DDRGK-motif had no effect on fertility, *allc1* is a male fertility gene, like *sun1* (Table S1). ALLC1 additionally pulled down a karyopherin alpha subunit, which may be essential^27^ and kinesin-15, which is specifically found in the cytosol of male gametocytes and first needed for the formation of motile gametes^62^.

**Figure 7.**
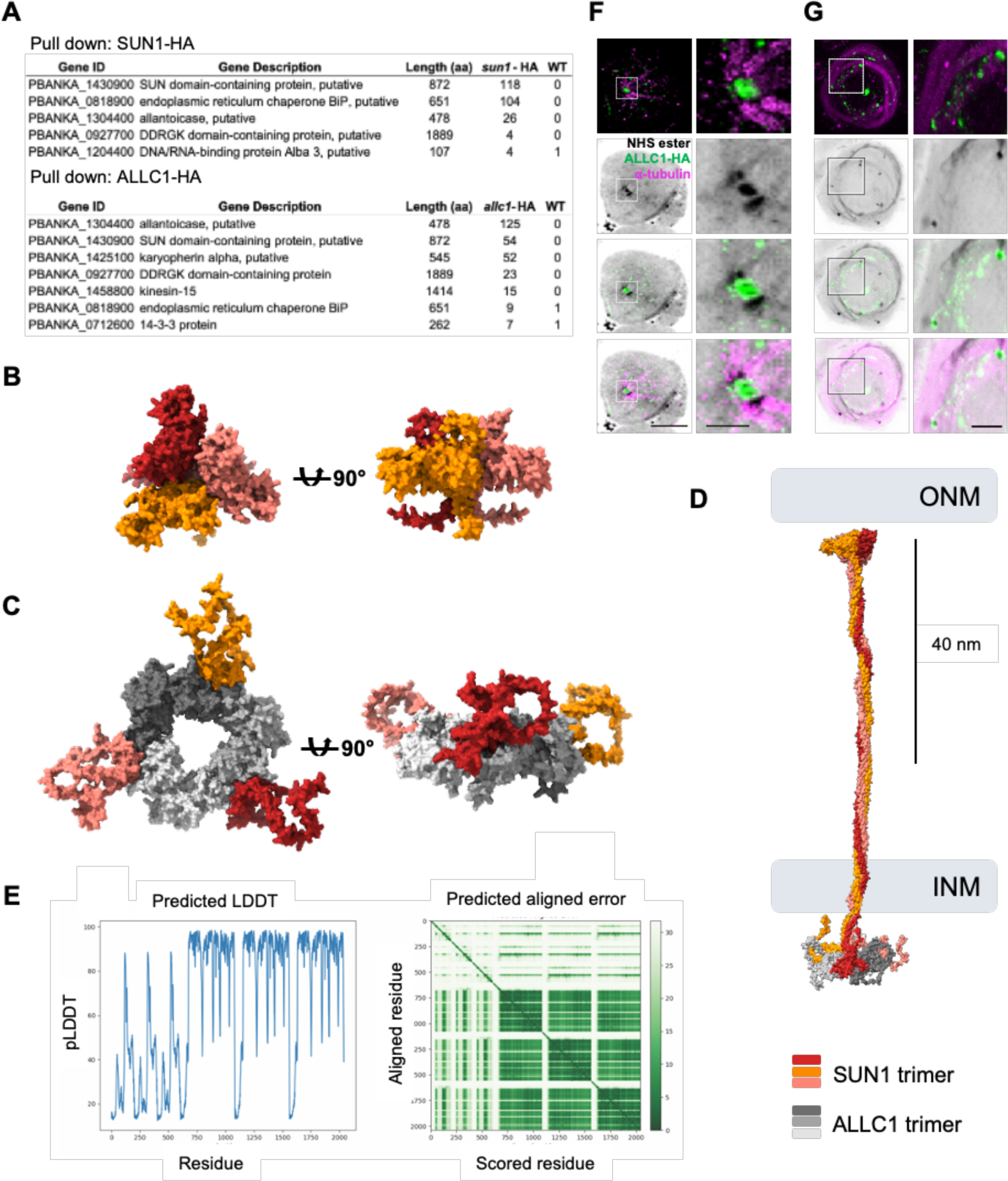
Identification and characterisation of a SUN1-interacting allantoicase-like protein, ALLC1. (A) Tables of putative SUN1 and ALLC1 interacting proteins identified by mass spectrometry analysis of proteins immunoprecipitated from activated gametocytes using HA-tagged targets. (B) AlphaFold model of a trimer composed of the C-termini of SUN1, containing the SUN domain. (C) AlphaFold model of an ALLC1 trimer (in shades of grey) in complex with the N-terminal domains of a SUN1 trimer. (D) AlphaFold model of the complete SUN1-ALLC1 complex. (E) Predicted LDDT and aligned error AlphaFold2 plots show the confidence of the interaction between SUN and ALLC1 residues. (F) Expansion microscopy of a microgametocyte fixed two minutes after activation, with immunodetection of ALLC1-HA and α-tubulin, and using Atto 594 NHS ester as general protein stain. Scale bars represent 5 μm of the expanded cell, 1 μm for inserts. (G) As above, except that cells were fixed eight minutes after activation.

Using AlphaFold2, we modelled a complex composed of three SUN1 proteins interacting with an ALLC1 trimer (Fig. 7B-D). The modelled complex is characterised by interacting C-terminal SUN domains, which in other eukaryotes would each bind to KASH (Klarsicht, Anc-1, Syne-1 homology) domain proteins of the outer nuclear membrane (Fig. 7B). We predict the N-terminal SUN1 domains to interact with an ALLC1 trimer (Fig. 7C), possibly replacing the nuclear lamins of the canonical LINC complex, which appear absent in *Plasmodium*. Connecting these two parts of the complex is a stretch of α-helical coiled coils, which may be as long as 80 nm (Fig. 7D, E). Consistent with the structural model, expansion fluorescence microscopy localises ALLC1-HA to the MTOCs of activated microgametocytes (Fig. 7F-G). Like with SUN1-HA, some of the cellular protein localised to the constriction between the nuclear and the cytosolic part of the MTOC.

The evolutionarily conserved C-terminal SUN domain folds into a compact β-sandwich connected to an N-terminally positioned α-helix to resemble a cloverleaf^63,64^. AlphaFold2 modelling of SUN1-ALLC1 suggests that a hetero-hexameric complex forms when homotrimeric ALLC1 binds the homotrimeric N-terminal region of SUN1 in the nucleoplasm. While crystallographic studies show that SUN domain proteins form stable homotrimers and bind KASH proteins in the perinuclear space via the C-terminal SUN domain^63,64^, the structural biology of protein-protein interactions in the nucleoplasm via the N-terminal region of SUN has not been described.

## DISCUSSION

Sexual reproduction and infection of the mosquito vector are tightly linked in malaria parasites and essential for disease transmission. The ploidy transition at fertilisation has so far posed an insurmountable challenge for the systematic identification of essential genes uniquely involved in both these processes. By mutagenising each sex separately with a library of barcoded gene targeting vectors, we have here overcome a major roadblock to revealing the biology of sex and transmission in a rodent malaria species. Through genome-scale genetic screens we have identified more than 400 fertility functions among targetable genes, increasing by more than an order of magnitude the number of transmission genes whose functions have been assayed for each sex individually in genetic crosses. We used the advantage of barcoded *Plasmo*GEM vectors of making bespoke vector pools to stratify male sterility genes further in secondary screens for egress and motility scores. Benchmarking our results against published phenotypes, we observe a good degree of consistency. We also find the new data recapitulate a smaller number of hits from previous screens for gametocyte development, which captured only one aspect of fertility^34^. There are limitations to this work. First, the *Plasmo*GEM project could generate barcoded targeting vectors for only 65% of the core genome, so the screens are blind to the remaining genes. Second, genes previously found essential for growth of asexual blood stages (>40% of genes) were excluded from the current experiments to give maximal space to those mutants expected to add to the gametocyte pool, and 388 slow growing mutants were tested separately, resulting in around 1300 genes that were tractable by our approach.

Aided by AlphaFold structure-based annotations, the new phenotype data provide deep insights into the biology underpinning both, malaria transmission and sexual reproduction in a divergent eukaryote. The data identify new genes involved in metabolic reprogramming of mitochondria, whose increased reliance on oxidative phosphorylation renders transmission particularly sensitive to drugs such as atovaquone. Among the most essential fertility genes, we found druggable enzymes of unknown function, such as a putative second lactate dehydrogenase essential specifically for male fertility (PBANKA_1340400) and a putative peptidyl-prolyl *cis-trans* isomerase (PBANKA_1143200), essential for fertility in both sexes, which does not affect expression of gametocyte reporter genes^34^ but where males have a low egress score, indicating a role late in gametocyte development. Our functional data can now also guide the discovery of essential surface-exposed proteins as new targets for transmission blocking vaccines, exemplified by the male gamete fusion protein *hap2*^9^, which may have a female counterpart.

In malaria parasites, some hallmarks of eukaryotic cells are uniquely important for reproduction and transmission. Starting are revealed for future studies elucidating such biological systems, including stage-specific metabolism and meiosis, a better understanding of which in *Plasmodium* will shed light on the evolution of an important aspect of eukaryotic biology. Similarly, our understanding of axoneme ultrastructure and function in apicomplexan parasites is rudimentary. Guided by structure homology and a phenotypic assay that tests a correlate of flagellar motility, we describe more than a dozen putative axoneme components with homologues among conserved eukaryotic flagellar proteins. Many of these were revealed only recently, through structural studies in human sperm or model organisms^65,66^ and some have not been studied experimentally in any species. More axoneme components may be discovered among the unannotated non-motile mutants from the screens. We propose that *P. berghei* can provide a genetically tractable model to study biomechanical aspects of ciliary motility and to develop ways to use such knowledge to block malaria transmission.

Studying conserved *Plasmodium* genes with fertility functions can reveal new insights into potentially ancient aspects of sexual biology. This is illustrated by our analysis of the nuclear envelope protein SUN1, which we show to be required for the MTOC to execute its nuclear functions during microgamete formation. The single bipartite MTOC, which in non-activated male gametocytes is already associated with the nuclear envelope, is composed of a cytosolic part marked by centrin, and an intranuclear body (INB)^14,67^. Upon activation, the cytosolic part of the MTOC gives rise to two ‘layered’ tetrads of centrioles, each of which assemble an axoneme^67^. Through three rounds of closed mitosis, MTOCs are then sequentially split up and moved by unknown forces into opposition along the nuclear envelope, with mitotic spindles emanating from their nuclear compartments. We find that SUN1 is not required for the cytosolic MTOC compartment to nucleate eight axonemes, which extend to a normal length and become motile. However, MTOCs in a *sun1* mutant do not divide, and axonemes remain connected through a single MTOC, which resides in the cytoplasm, distant from the nuclear envelope, and no mitotic spindles form. Individual microgametes can probably break away from the MTOC when their axonemes start beating, but consistent with the failure of the MTOC to insert into the nuclear envelope and the lack of spindles, free microgametes are mostly anucleate.

We propose that the recruitment or insertion of the microgametocyte’s MTOC into the nuclear envelope either rely directly on SUN1, or that SUN1 is required for the formation of mitotic spindles, which may in turn keep the MTOC anchored indirectly in the nuclear envelope. Either way, SUN1 is critical for linking the male genome to the gamete’s axoneme. In contrast, neither the rapid threefold replication of the MTOCs upon gametocyte activation, nor the replication of the genome rely on SUN1 and therefore on an intact mitotic machinery, underlining the paucity of cell cycle checkpoints in *Plasmodium*. The anucleate microgametes of the *sun1* mutant are only superficially similar to a mutant lacking the microtubule end binding protein EB1, which in *Plasmodium* has a non-canonical function of binding laterally to microtubules, thereby linking kinetochores to spindle microtubules^68^. Both *eb1* and *sun1* mutants have anucleate microgametes, but for different reasons -- while in the *eb1* KO mitotic spindles fail to take along chromosomes, in the *sun1* KO, axonemes and MTOCs are dissociated from nuclei, and spindles do not form.

When combined with evidence from model eukaryotes, our data support an ancient role of a SUN domain protein in the formation of flagellated gametes: (1) In *S. pombe*, which also perform a closed mitosis, the SUN protein Sad1 localises to the functional equivalent of intranuclear body of the *Plasmodium* MTOC, the spindle pole body (SPB), where it is needed for mitotic spindles to form and possibly for embedding the SPB in the nuclear envelope^69^. (2) In mice, sperm associated antigen 4 (Spag4 = SUN4) is a spermatid nuclear envelope protein with a Sad1-like C-terminal SUN domain critical for sperm head formation and fertility^70,71^. (3) Similarly, in *Drosophila* spermatids, a Spag4 homologue is a fertility factor that anchors the basal body to the spermatid nucleus through a mechanism which, like in *Plasmodium*, does not appear to involve a canonical LINC complex^72^.

We find that *P. berghei* SUN1 forms a complex with another male fertility protein, ALLC1. The interaction is well supported by reciprocal immunoprecipitation, co-expression, similar phenotypes in our fertility and motility screens and localisation to pore-like structures surrounding the base of the intranuclear body of the MTOC at the nuclear envelope. Structural modelling supports a direct interaction between an ALLC1 trimer and the N-terminal part of a SUN1 trimer, where canonical SUN proteins would interact with lamins in the nucleoplasmic periphery. In the interacting region of SUN1, serine 43 is dephosphosphorylated upon gametocyte activation^73^, which may be significant because in *C. elegans* SUN1 the homologous residue, S273, is required for a non-lamin interactor to localise centrosomes to the nuclear envelope during embryogenesis^74^. SUN domain proteins typically reside in the inner membrane of the nuclear envelope, and if their typical topology^75^ is conserved in a SUN1-ALLC1 complex lining the nuclear pore, both the N-terminus of SUN1 and the ALLC1 subunits of the complex would be positioned to interact with spindle microtubules, possibly through the kinesin-15 homolog that co-precipitates exclusively with ALLC1. *P. berghei* has a second putative allantoicase, PBANKA_0213200, which forms part of the kinetochore complex in the microgametocyte^76^, intriguingly placing two different allantoicase at each end of the mitotic spindle.

ALLC1 is unlikely to function in purine degradation in *Plasmodium* since current models of metabolism do not suggest this pathway exists^17,77^. Similarly, vertebrates lack allantoicase activity, although homologous genes are present. Intriguingly, mouse allantoicase transcript abundance is highest in testis, and the protein has been suggested a male contraceptive target until shown to be dispensable for male fertility^78^. Our data thus not only provide concrete evidence for an alternative function of an allantoicase, but also raise the intriguing possibility that such a function may be conserved in eukaryotes where SUN1 domain proteins perform mitotic or fertility functions outside of the canonical LINC complex. Altogether, our data provide a broad and unbiased picture of the molecular processes that underpin malaria parasite transmission but also highlight ancestral aspects of sex that have evolved close to the last eukaryotic common ancestor. Conserved structures include the axoneme and meiotic machinery and may extend to an evolutionary connection between SUN domain proteins functioning outside of the canonical LINC complex to organise MTOCs in *Plasmodium* gametogenesis, in the yeast spindle pole body and vertebrate sperm.

## METHODS

### Parasites and mosquitoes

All experiments used the reference clone cl15cy1 of *P. berghei* ANKA. *Anopheles stephensi* mosquitoes were bred in-house at 28 °C and 80 % humidity. Following infection with *P. berghei* parasites, mosquitoes were maintained at 19 °C and 80 % humidity. All mosquitoes were supplied with 8 % fructose solution (filter-sterilised, supplemented with PABA), and anaesthetised with CO^2^ gas and then on ice for dissection.

### Rodents

Rodent research at the Wellcome Sanger Institute was conducted under licence from the UK Home Office, and protocols were approved by the Animal Welfare and Ethical Review Body of the Wellcome Sanger Institute. Rodents were kept in pathogen-free conditions and subjected to regular pathogen monitoring by sentinel screening. They were housed in individually ventilated cages with autoclaved aspen wood chips, a fun tunnel and Nestlets at 21 ± 2 °C under a 12:12 hour light-dark cycle at a relative humidity of 55 ± 10 %. They were fed a commercially prepared autoclaved dry rodent diet and water, both available ad libitum. The health of the animals was monitored by routine daily visual health checks. The parasitaemia of infected animals was determined by methanol-fixed and Giemsa-stained blood smears. Female RCC Han Wistar outbred rats (Envigo, UK) aged eight to 14 weeks were infected with *P. berghei* WT parasites by intraperitoneal injection. Infected rats served as donors for *ex vivo* schizont cultures on day five of infection, at a parasitaemia of 1-5 %. Rats were housed with one cage companion. Rats were terminally anaesthetised by vapourised isoflurane administered by inhalation prior to terminal bleed. Rats were used because they give rise to more schizonts with higher transfection efficiency compared to mice. Transfection efficiency is critical when screening pools of vectors. Transfected parasites were injected intravenously into the tail of female adult BALB/c inbred mice aged 8-22 weeks (median age 10 weeks). This animal model was chosen to minimise host genetic variability and to obtain robust infections with a low incidence of cerebral pathology.

Animal research at Umeå University was conducted under Ethics Permit A13-2019 and approved by the Swedish Board of Agriculture. Mice were group-housed as four cage companions, and rats as two cage companions, in IVC with autoclaved wood chips and paper towels for nesting, at 21 ± 1°C under a 12:12 hr light dark cycle at a relative humidity of 55 % ± 5 %. Specific-pathogen-free conditions were maintained and subjected to Exhaust Air Dust (EAD) monitoring and analysis biannually. Mice and rats used at Umeå University were purchased from Charles River Europe. Female BALB/c mice aged 6-20 weeks old were used, and female Wistar Han IGS rats were ordered at 150 g, and used within two months.

## METHOD DETAILS

### Parasite maintenance and life cycle analysis

The *P. berghei* strain ANKA cl15cy1 reference line was used for WT control parasites and transfection background lines. Parasites in fresh or frozen blood stocks were used to infect mice or rats by intraperitoneal injection, and parasites after transfection or FACS were used to infect mice by tail intravenous injection. Parasites were harvested from rodents by cardiac puncture. When high gametocytaemia was required, mice were treated with 200 ul phenylhydrazine (6 mg/ml; Sigma) by tail intravenous injection two to three days before infection. For exflagellation, parasites were activated on day three to five post-infection at 19-21 °C in approximately five volumes of exflagellation medium (RPMI medium 1640, 100 µM xanthurenic acid, pH 8.2) or ookinete medium (RPMI medium 1640, 100 µM xanthurenic acid, sodium bicarbonate, 100 U/ml penicillin-streptomycin, 20 % v/v heat-inactivated FBS). Exflagellation events were quantified by light microscopy, and a haemocytometer was used when the number of exflagellation events per 10^5^ erythrocytes was required. To generate ookinetes *in vitro*, heavily infected mice were cardiac bled into 10-20 volumes of ookinete medium on day four to five post-infection, which was incubated at 19 °C for 20-24 hours. To determine ookinete conversion rates, and quantify DNA content, P28-positive macrogametes and ookinetes were counted and imaged by fluorescent microscopy after incubating with 1 mg/ml Cy3-conjugated 13.1 monoclonal anti-P28 antibody^79^ and Hoechst 33342 (1 mg/ml). To assess oocyst production, on day three post-infection, mice were anaesthetised and placed on mosquito cages. Starved mosquitoes were allowed to blood feed for 15 minutes. On day 10-11 post-mosquito infection, 20-30 mosquito midguts were dissected, and oocysts were counted by phase-contrast microscopy.

### Single-sex background lines

#### Female-only line

The *Plasmo*GEM Gene Out Marker Out (GOMO) transfection vector (*Pb*GEM-646728) targeting the male development 4 gene (*md4*; PBANKA_0102400), was transfected into the *Pb*ANKA cl15cy1 reference line and selected by pyrimethamine (0.07 mg/ml; MP Biomedicals). Before parasitaemia reached 1 %, 30-50 parasites expressing both GFP and mCherry were obtained by FACS and intravenously injected into the tail of naive mice. Once parasites were observed, the mice were switched to 5-fluorocytosine (1 mg/ml; Sigma) drinking water to negatively select against parasites that express mCherry, and therefore MD4. Before parasiteamia reached 1 %, FACS was used to obtain 30-40 parasites that expressed GFP only, which were intravenously injected into the tail of naive mice. The GFP-only parasites were challenged with pyrimethamine, which confirmed that they were marker-free. Additionally, PCR genotyping confirmed that the parasites do not contain DHFR or mCherry (Table S5; Figure S1).

#### Male-only line

As there was no *fd1* GOMO transfection vector available from *Plasmo*GEM at the time, we constructed a GOMO transfection vector to target *fd1* (PBANKA_1454800). 5’ and 3’ homologous gene flanking regions with *Not*I sites and Gibson tails were amplified by PCR (Table S5). A pCC1 vector was digested with *Sac*II/*EcoRI*, and an R6K-R1R2_ZP vector was digested with *Not*I/*Avr*II. The ampicillin resistance-containing fragment was purified from the pCC1 digestion, but the zeoPheS-containing fragment was not purified from the R6K-R1R2_ZP digestion, due to a very small size difference. A Gibson assembly was performed with the 5’ and 3’ homologous gene flanking regions, the purified pCC1 fragment and the purified R6K-R1R2_ZP digestion, which was transformed into XL10-Gold cells. Colonies were PCR-screened for the presence of the 5’ and 3’ flanks (Table S5), and then checked by *Spe*I/*Bam*HI digestion and capillary sequencing (Table S5). Correct vectors were added to a Gateway reaction with the GW21-R6K-GOMO-GFP-Cherry plasmid, which was transformed and analysed by PCR-screening, *Not*I/*Bam*HI digestion, and capillary sequencing (Table S5). A verified transfection vector was transfected into the *Pb*ANKA cl15cy1 parasite line and made marker-free as above. Genotyping PCRs confirmed correct integration of the transfection vector, and absence of DHFR and mCherry (Table S5; Fig. S3).

### Pilot and genome-wide fertility screens

#### Generation of mutant parasite pools

To determine mutant fertility by barcode sequencing (barseq), *Plasmo*GEM KO vectors (see Table S5 for all vector IDs) were pooled as shown in Table S1. Barcodes were counted by sequencing amplicons from the genomic DNA extracted from 50 midguts per replicate, each infected with hundreds of oocysts. Despite the high transmission efficiency of *P. berghei*, only 25,000-50,000 oocysts can be produced from an infected mouse, which sets a limit for the complexity of pools that can be sampled in a representative manner. In preliminary experiments we determined that pools of 100-200 mutants were optimal for measuring male phenotypes, but that to quantify female phenotypes with the same statistical power, only around 50 mutants could be transmitted together. Throughout the screens, we assessed biological variance by transmitting each pool at least twice from different infected mice. We further monitored sampling errors by subsampling each transmission experiment in two batches of mosquitoes and by amplifying and counting barcodes twice from each DNA sample. Data from all types of replicates were entered into the final analysis and used to determine independently for each sex the confidence with which a mutant’s transmission rate was determined. Each transmitted pool included the mutants with normal asexual blood-stage growth and fertility to serve as references against which the fertility of all other mutants was calculated, allowing data from nearly two hundred transmission experiments and >3000 individually dissected midguts to be combined into two datasets for male and female fertility.

The maximal pool sizes and required number of replicates were determined by monitoring the reproducibility and variance in a pilot screen of 58 vectors pooled for duplicate transfections into WT, female and male lines. In subsequent experiments, 43-202 and 193-204 vectors were prepared for duplicate transfections into female and male lines respectively. Smaller pools were made to improve representation of mutants in the female line, which appeared to have a lower intrinsic transfection efficiency. Vectors to generate reference mutants (approximately 100 ng per vector per transfection) were added to the pool DNA (approximately 10 µg per transfection), and the combined vectors were digested overnight with *Not*I to release the gene-targeting vectors from the vector backbones. Mutants with known dispensable phenotypes served as WT controls for normalising mutant barcode counts in each pool. Digested DNA was precipitated with ethanol and resuspended in nuclease-free water (7 µl per transfection). WT, female and male line schizonts from rats were harvested from 20-22 hour cultures with MACS Cell Separation Columns (Miltenyi Biotec). For each transfection, 4 µl of isolated schizonts was added to 7 µl of DNA and 18 µl of P3 Primary Cell 4D-Nucleofector solution (Lonza), of which 26 µl was added to a well in a 4D-Nucleofector 16-well strip (Lonza) and electroporated with the FI-115 program. Transfected schizonts were intravenously injected into the tail of naive mice and selected by adding pyrimethamine to the drinking water (0.07 mg/ml) from day one post-transfection. Residual liquid was collected from each well to determine the composition of the transfected vector pool.

#### Transmission of mutant parasite pools

On day three after transfection, naive mice were infected with both single-sex lines, and on day seven, blood from all infected mice was collected. To enable transmission, each blood containing a mutagenised sex was mixed with blood containing wild type parasites producing the complementary sex at a one-to-one ratio based on parasitaemia. Three naive mice were infected with each mixture for each of two duplicate transmission experiments. Infected mice were anaesthetised and used to feed approximately 400 female mosquitoes. For each transmission experiment, blood from each of the three mice was collected and pooled for the blood input sample. On day 13 or 14 post-mosquito infection, approximately 50 infected midguts were collected in duplicate from the same cage of mosquitoes for two oocyst output samples per mosquito feed.

#### DNA extraction and Illumina sequencing

KO vector pool DNA was extracted from the well input samples in PBS by incubating at 95 °C for 5 minutes. Parasite genomic DNA (gDNA) was extracted from a total of 21 µl of the blood input sample, with the DNeasy Blood and Tissue Kit (Qiagen), and eluted in 50 µl. Midgut oocyst output gDNA was extracted as described^75^. Briefly, 100 µl of QuickExtract DNA Extraction Solution (Epicentre) was added to a sample containing approximately 50 infected midguts, which was incubated at 65 °C with shaking (300 rpm) for six minutes, then 98 °C for two minutes. Mutant barcodes were amplified from the DNA samples in duplicate using a nested PCR^27^, before sample-specific index tags were added to each sample so they could be pooled in groups of approximately 36. The libraries were sequenced using the Illumina MiSeq Reagent Kit v2 (300 cycles), which were loaded at a low cluster density (4x10^5^ clusters/mm^2^) with 50 % PhiX.

### Male motility subscreens

#### Generation of male mutant parasite pools

To screen the top 125 male mutants in dispensable genes^27^ for egress and flagellar motility we created, digested and precipitated two vector pools spiked-in control vectors. These were transfected into schizonts from the male-only line in duplicate, and replicates from the different pools were combined prior to injection into the tails of two naive mice, so that each mouse received 125 mutants that were selected by pyrimethamine from day one post-transfection.

#### Secondary screens for egress and motility

On day seven post-transfection, pyrimethamine-selected parasites were harvested and passaged into phenylhydrazine-treated mice to increase gametocytaemia. On day three, asexual parasites were killed by intraperitoneal injection of sulphadiazine (0.3 mg/kg). After 24 hours, infected blood containing predominantly gametocytes was collected for microgamete purification as described^54^. Briefly, parasites were activated for 20 minutes and subjected to a two-step centrifugation protocol. In the first step egressed gametes were retained in the supernatant due to their lower density. In the second step all egressed parasites were pelleted, and motile parasites were allowed to swim for 10 minutes into the supernatant which was then separated from the cell pellet. DNA was extracted from all fractions and barcodes were counted to ask which mutants were blocked before egress, and which of those that egressed became motile enough to swim back up into the supernatant after sedimentation. DNA libraries were prepared and sequenced as described above.

### Phenotypic analysis of a male SUN protein

#### Single transfections and genotyping

The *Plasmo*GEM PBANKA_1430900 (SUN1) KO and HA-tagging transfection vectors (Table S5; Fig. S3) were individually transfected into the *Pb*ANKA Cl15 cy1 parasite background line. Pyrimethamine-selected parasites were genotyped by PCR (Table S5; Fig. S3), which confirmed integration of the transfection vectors and little WT contamination. To ensure that any phenotype observed is from a KO parasite, *sun1* KO parasite sub-clones were obtained by dilution cloning (Table S5; Fig. S3).

#### Western blotting

To ensure that the tagged parasites express HA, saponin-isolated SUN-HA parasites from a heavily infected phenylhydrazine-treated mouse harvested on day three post-infection, were resuspended in reducing buffer and heated to 70 °C before analysis. The protein samples were electrophoresed using the Mini-PROTEAN Electrophoresis System (BioRad) and transferred to a polyvinyl difluoride membrane (BioRad). After blocking, membranes were incubated with a monoclonal rabbit anti-HA antibody (1:1000; Cell Signaling Technologies). After washing, membranes were incubated with an anti-rabbit HRP antibody (1:20,000; Cell Signaling Technologies). Protein bands were detected with the Immobilon Western Chemiluminescent HRP Substrate (Millipore).

#### DNA content analysis

Microgametocytes and ookinetes were fixed and DNA was stained with Hoechst. Hoechst images were captured for microgametocytes, ookinetes, rings and merozoites. Microgamete, ookinete, 1 N control cell, and background Hoechst intensity was measured in Fiji, and the DNA content of microgametes and ookinetes was determined by subtracting the background and normalising to 1 N control cells.

#### Immunofluorescence assays

Highly infected phenylhydrazine-treated mice were bled on day four-five post-infection. The blood was immediately added to exflagellation medium at 19 °C. At the appropriate time points, aliquots were added to paraformaldehyde in microtubule-stabilising buffer (MTSB; 10 mM MES, 150 mM NaCl, 5 mM EGTA, 5 mM MgCl^2^, 5 mM glucose, pH 6.9) with a final concentration of 4 % paraformaldehyde and fixed at room temperature for at least 30 mins. Fixed cells were washed with MTSB and stored at 4 °C overnight. Fixed cells were loaded at a concentration of approximately 1 % onto poly-D-lysine-coated (Sigma) slides and incubated at room temperature for 15 minutes. Cells were permeabilised with 0.2 % TritonX-100 and blocked with 3 % BSA. Cells were incubated in primary antibodies in 3 % BSA, washed, then incubated in secondary antibodies in 3 % BSA. Hoechst (1:1000) was included in the final wash, and samples were finally mounted in Vectashield (Vector laboratories).

#### Immunoprecipitation and mass spectrometry

WT, SUN1-HA and ALLC1-HA parasites at eight minutes post-activation were used to prepare cell lysates. Purified parasite pellets were homogenised in IPP150 lysis buffer (50 mM Tris-HCl pH 8, 150 mM NaCl, 1 % NP-40, 1 mM EDTA, 1 mM DTT and protease inhibitors). Lysates were centrifuged at 20,000 g for 15 minutes at 4 °C. Immunoprecipitations were performed by incubating the lysate supernatants with Protein-G Dynabeads (Invitrogen) coupled with an anti-HA antibody (Roche), for two hours with shaking at 4 °C. To prepare for LC-MS/MS, the samples were washed with cold IPP150 buffer then cold 50 mM ammonium bicarbonate. The bound proteins were digested with trypsin overnight and the protein elutions were collected and added to 50 mM ammonium carbonate. Peptides were analysed by mass spectrometry and raw data processed as described^34^.

#### Ultrastructure expansion microscopy

Parasites were generated, activated and fixed as above. Fixed cells were added to poly-D-lysine-coated (Gibco) coverslips at an approximate cell concentration of 2 % on the day of fixation and incubated for 15 minutes. Protein crosslinking prevention, gelation, denaturation and gel expansion were done as described^60^. Gels were incubated in rabbit anti-HA (1:250; Cell Signaling Technology), and mouse anti-α-tubulin (1:250; Sigma) or mouse anti-centrin (1:500; Sigma) in 2 % BSA at 37 °C for three hours. Primary antibodies were discarded, and gels were washed with PBS-Tween 0.1 % before incubating in goat anti-rabbit Alexa Fluor 488 (1:400, Thermo Fisher) and goat anti-mouse Alexa Fluor 405 (1:400; Thermo Fisher) at 37 °C for 2.5 hours. Secondary antibodies were removed, and gels were washed as before. Cells were stained with Atto 594 NHS Ester (10 µg/ml; Merck) in 1X PBS on an orbital shaker at room temperature for 1.5 hours. The stain was discarded, and gels were washed as before. Gels were expanded again overnight and mounted and imaged as described^60^. Confocal images were acquired on a Leica SP8 Confocal microscope (HC PL APO 63x/1.40 OIL CS2; Diode 405nm, Diode 638 nm, OPSL 488 nm, OPSL 552nm), and widefield images were acquired on a Zeiss Axio Imager 2 microscope. All images were analysed and processed in Fiji^80^.

## QUANTIFICATION AND STATISTICAL ANALYSIS

### Analysing barseq data

For each pool of mutants, index tags were used to separate raw sequencing reads into each sample, and a python script counted the number of mutant barcode sequences^25,27,75^. The relative abundance of a mutant was calculated by dividing the mutant barcode count by the total number of mutant barcodes in a sample. The standard deviation was computed for PCR duplicate samples to assess the technical variation.

### Removing noise from barseq data

Mutants with relative abundances of zero in KO vector pool input samples were considered technical dropouts and removed from subsequent analyses. For remaining mutants, one was added to each relative abundance value to avoid infinity in the log^2^ transformation. To ensure confidence in our analysis, we only assigned phenotypes if the log^2^ relative abundance in the relevant input sample (fertility screen blood or motility screen pellet) was above the threshold for barcode count noise. To determine the threshold, we plotted input log^2^ relative abundance against relative error (standard deviation normalised to relative abundance). This allowed us to identify real mutant relative abundances, which were above log^2^ -12 in fertility screen blood input samples. Mutants below this threshold in the fertility screen were assigned ‘insufficient data’. All mutant relative abundances in the motility screen pellet input samples could be analysed with confidence.

### Calculating relative fertility or motility

To determine the relative fertility of each mutant in each fertility screen pool, and the relative motility of each male mutant in the motility screen pool, the relevant input sample log2 relative abundance was subtracted from the relevant output sample log2 relative abundance, which was then normalised to WT controls. Error values were calculated for each mutant by propagating the standard deviations associated with each sample. Multiple relative fertility or motility values for each mutant were ultimately combined using the inverse-variance weighted mean.

### Assigning phenotypes

The difference of maximum (diffmax) and minimum (diffmin) were used to assign phenotypes for both screens, where diffmax or diffmin are equal to the relative fertility or relative motility value plus two standard deviations. To define the fertility phenotype of mutants in the fertility screen, mutants with a diffmax less than -2 were called ‘reduced’, and mutants with a diffmax greater than -2 were called ‘not reduced’. To define the motility phenotype of mutants in the male motility screen, mutants with a diffmin greater than -1 were called ‘not reduced’, and mutants with a diffmin less than -1 were called ‘reduced’.

### Enrichment analysis of sex-specific fertility genes

Sex-specific fertility genes were analysed in PlasmoDB for gene ontology enrichment, including biological process, cellular component and molecular function. Significant enrichment of sex-specific fertility genes in MPMP pathways was also assessed using the hypergeometric probability density function (P), where k and K are the numbers of sex-specific fertility genes and the number of total genes in a pathway, respectively, and n and N are the total number of sex-specific fertility genes in the screen and the total number of genes screened, respectively (equation 1). For each pathway, the p-value was calculated by the sum over probabilities for greater or equal than the enriched number of sex-specific fertility genes to test the null hypothesis (equation 2).

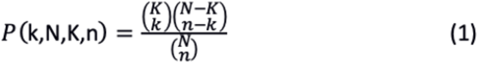

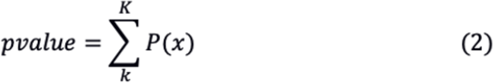

### Functional annotation via structural modelling

The overall workflow was as described^81^. Briefly, we folded the proteins indicated to be important in the genetic screen using ColabFold with default parameters. Next, we used each individual predicted 3D structure as input for Foldseek searches employing the alignment type 3Di+AA Gotoh-Smith-Waterman (local, default) and ran it against three different databases: 1. PDB, 2. AlphaFold database from the 20 first annotated model organisms (accession date: 07-15-2022), and 3. SwissProt AlphaFold. Predicted models, and their structural homologs were visualised in our Chimera X plugin, to allow for manual curation and the functional annotation.

### Structural modelling of the SUN1-ALLC1 complex

Sequences for SUN and ALLC proteins from *Pb*ANKA were identified by BLAST searching (tBLASTn) and searching Uniprot (https://www.uniprot.org) for related gene annotations. Sequences were assessed for relationship by BLAST analysis, Clustal Omega alignments (https://www.ebi.ac.uk/Tools/msa/clustalo/), structural comparisons by AlphaFold monomer alignments and reverse searching related structures using Foldseek (https://search.foldseek.com/search)^82–85^.

SUN protein sequences were trimmed from the first predicted membrane-spanning domain as annotated on UniProt. The predicted inner nuclear component was run on AlphaFold2 v2.2 or 2.3 in multimer mode on custom Cloud instances hosted by AWS configured with the default installation of AlphaFold as described at the Github site (https://github.com/google-deepmind/alphafold) or on a Google Cloud vertex installation custom engineered in collaboration with UNSW, Intersect, Google Australia and Deepmind. All instances were run using the default setting including using the full database with either five or 25 models and running Amber relaxation. The SUN1 protein and predicted ALLC1 protein from the same species and strain were run in multimer mode.

The best ranked models were compared structurally by structural alignment in Pymol v2.5.5 of the allantoicase domain and their pLDDT and PAE confidence regions were assessed for the SUN1-ALLC1 complex. Regions of the SUN1 protein that did not make contact with the ALLC1 trimer were removed from the model to facilitate identification of the interaction site and residues involved and were used in figure production.

## DATA AND SOFTWARE AVAILABILITY

https://github.com/vpandey-om/Fertility_screen http://obilab.molbiol.umu.se/pgem/phenotype

## AUTHOR CONTRIBUTIONS

Conceptualisation, O.B.; Methodology, O.B. and C.S.; Software, V.P.; Formal Analysis, O.B, C.S., V.P., M.P.C., D.S.; Investigation, C.S., A.B., M.H., M.P.C, K.M., D.S., O.B., V.P.; Writing – Original Draft, C.S.; Writing – Review & Editing, O.B., C.S., V.P.; Visualization, O.B., C.S., V.P., M.K.; Supervision, O.B., C.S., J.C., R.B.; Project Administration, C.S.

## Supporting information

Table S1 Fertility phenotypes

Table S2 Selected gene functions

Table S3 Motility and egress scores

Table S4 Protein pull down data

Table S5 Vectors plasmids and olives

## ACKNOWLEDGEMENTS

The authors are grateful to Tom Metcalf, Ondine Duverger, Stuart Wales, Sarah Lundgren and Rashmi Mishra, who bred and prepared the mosquitoes for this work, and to the *Plasmo*GEM team at the Wellcome Sanger Institute for making the vectors. Mandy Sanders and the sequencing team at the Wellcome Sanger Institute and the SciLifeLab National Genomics Infrastructure in Stockholm are acknowledged for sequencing support, as is the Structural Biology Facility within the Mark Wainwright Analytical Centre UNSW for structure analysis support.

## Funding

Work at Umeå University received funding from the Knut and Alice Wallenberg Foundation and the European Research Council (Grant agreement No. 788516). MH was supported by SNF (P2SKP3_187635), HFSP (LT000131/2020-L) and a Marie Sklodowska-Curie Action fellowship (No. 895744). Work at the Wellcome Sanger Institute was funded by Wellcome core grant 206194/Z/17/Z.

## Notes

### Competing Interest Statement

The authors have declared no competing interest.

