## Supplementary material for "Systematic screens for fertility genes essential for malaria parasite transmission reveal conserved aspects of sex in a divergent eukaryote": Figure S1

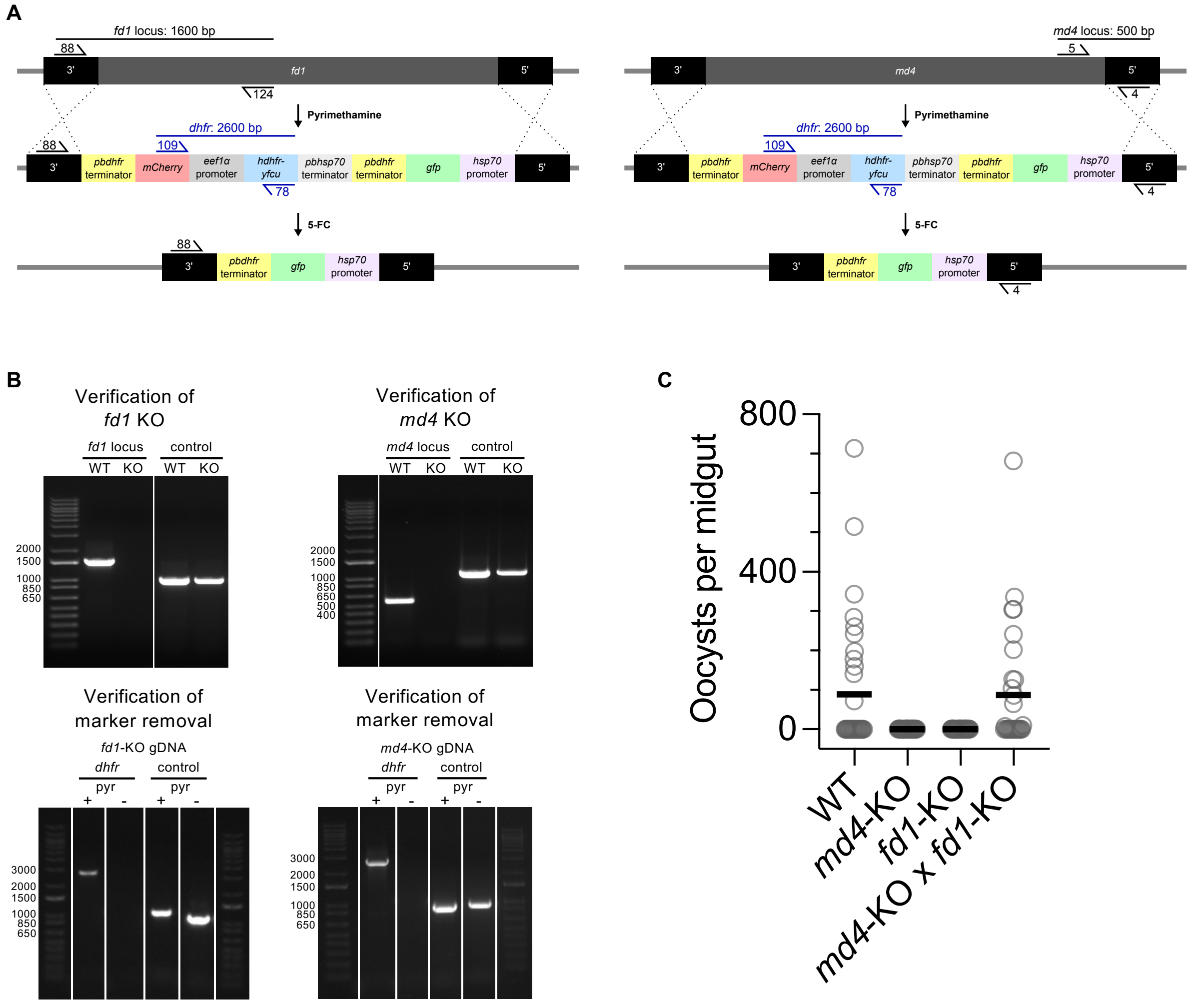


**Figure S1. Generation and characterisation of single-sex marker-free lines.**

A. Schematics showing the integration of the *Plasmo*GEM vectors for either *female development 1* (*fd1,* PBANKA_1454800) or *male development 4* genes (*md4*, PBANKA_0102400) leading to loss of female or male gametocytes, respectively (Russel *et al*., 2023) under pyrimethamine selection for *hdhfr*, followed by negative selection on 5-fluorocytosine (5-FC) against the *yfcu* gene to select for random recombinants between the directly repeated *pbdhfr* terminator sequences leading to loss of the *hdhfr-yfcu* selection cassette.

B. Genotyping PCR amplicons separated on ethidium bromide-stained agarose gels using primers shown in (A). Absence of intact *fd1* and *md4* loci was tested using primer pairs 88-125 and 5-4, respectively. A primer pair amplifying *pol2* served as positive control. Marker removal was tested with primer pair 109-78 on gDNA extracts obtained before ('+' with pyrimethamine) and after ('-' without drug) negative selection with 5-FC. See Table S5 for primer sequences.

C. Oocyst numbers on individual midguts 11-12 days after feeding with wild type and cloned knockout lines, as well as with a cross between the two mutant clones, initiated by coinfecting a mouse with both clones simultaneously.
