## Supplementary material for "Systematic screens for fertility genes essential for malaria parasite transmission reveal conserved aspects of sex in a divergent eukaryote": Figure S2

A

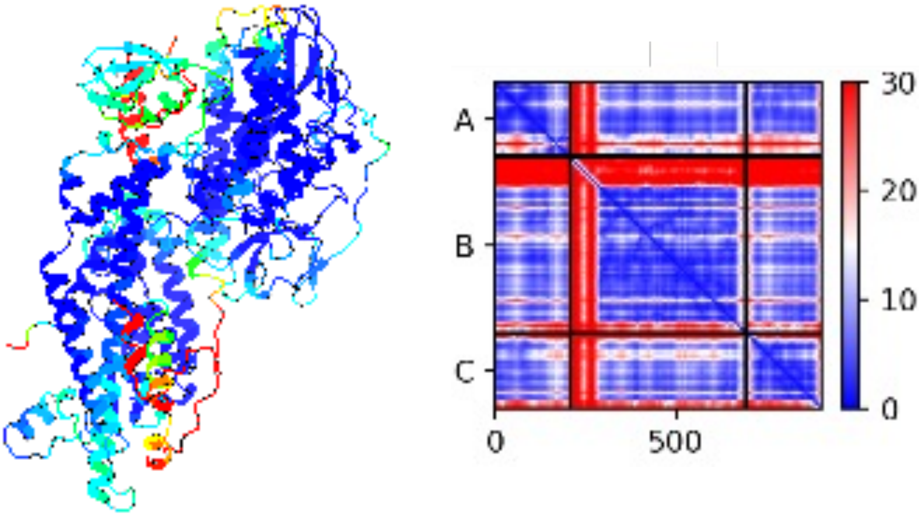


B

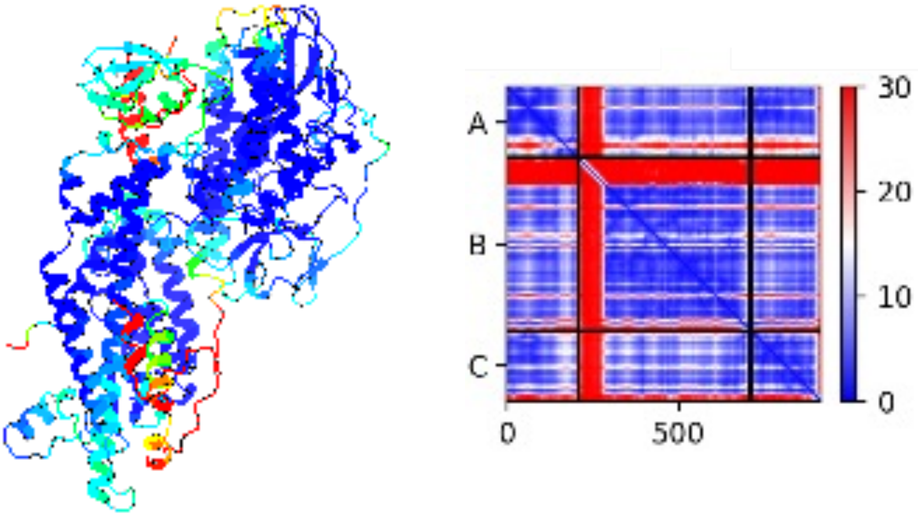


C

| *Homo sapiens* | *P. berghei* gene ID | *P. falciparum* asexual phenotype | *P. berghei* phenotype | Expression cluster (Cell atlas) |
|---|---|---|---|---|
| Sld5 + Psf1 | PBANKA_1238400 | Essential | n/d | MCA 11 |
| Psf2 | PBANKA_0807800 | Essential | n/d | MCA 8 |
| Psf2 | PBANKA_1349400 | Dispensable | Male fertility | MCA 12 |
| Psf3 | PBANKA_0815800 | Essential | n/d | MCA 11 |
| Psf3 | PBANKA_1402400 | Dispensable | Male fertility | MCA 12 |

**Figure S4. Duplication of GINS complex components in *Plasmodium*.**
(A, B) Structural models of the generic GINS complex (A) and the gametocyte-specific complex (B), coloured according to the pLDDT score from Alphafold2 (dark blue = high confidence, red = low confidence). The corresponding pae graph for the model is shown to the right.
(C) Table of GINS subunits showing duplication of Psf2 and Pfs3 in *Plasmodium*, phenotype data from Zhang et al. (2018) and the current study, and expression clusters according to Howick et al. (2019).
