## Supplementary material for "Systematic screens for fertility genes essential for malaria parasite transmission reveal conserved aspects of sex in a divergent eukaryote": Figures S3

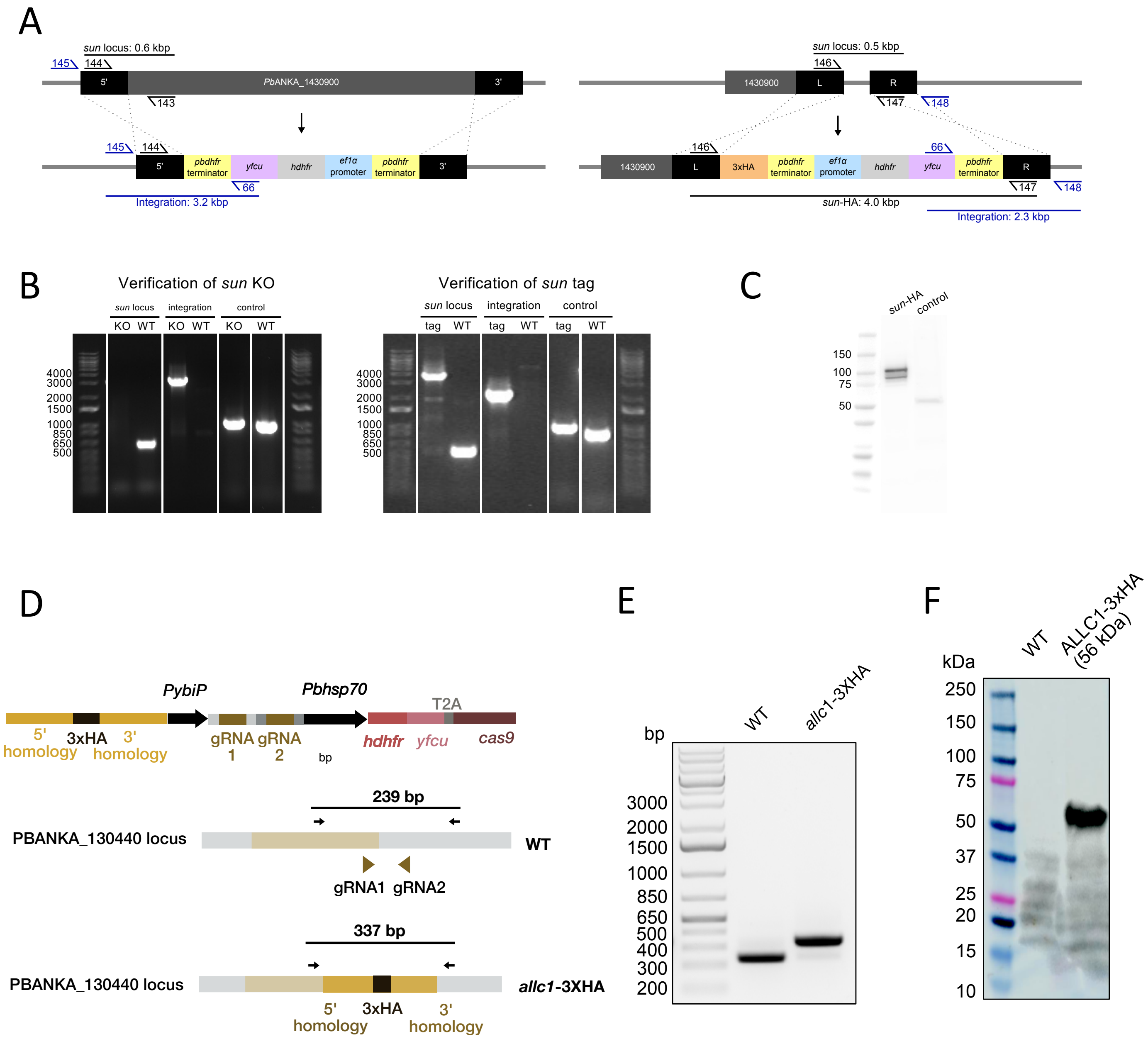


**Figure S3. Generation of *sun1*-KO and *sun1*-HA and *allc1*-HA lines.**

A. Schematics illustrating genomic integration of the *sun1* (*Pb*ANKA_1430900) KO and 3xHA-tag vectors.

B. Genotyping PCR amplicons separated on ethidium bromide-stained agarose gels using primers shown in panel (A).

C. Absence of the intact *sun1* locus was tested using primer pair 144-143 and correct integration of the KO vector was tested using primer pair 145-66. Presence of the HA-tagged *sun1* locus was tested using primer pair 146-147 and correct integration of the 3xHA-tag vector was tested using primer pair 148-66. A primer pair amplifying RNA polymerase II served as positive control with the same gDNA extracts.

D. Western blot of *sun*-HA parasite protein probed with rabbit anti-HA and anti-rabbit HRP antibodies. *rsph9*-HA parasite protein served as a negative control.

E. Schematic of the region with Cas9 plasmid used to generate C-terminally tagged allantoicase expressing lines and genomic locus of wild type ( WT) and *allc1*-3X HA tagged parasites. Guide RNA ( gRNA ) binding sites for primers are shown.

F. Genotyping PCR and western blot confirming the integration of 3xHA tag in *allc1* locus. All primer sequences are shown in Table S5.
